# A comparative study of genital lichen sclerosus transcriptomes

**DOI:** 10.64898/2025.12.10.692472

**Authors:** S. D. Margasyuk, A. L. Kuznetsova, D. A. Skvortsov, E. M. Alekberov, M. M. Iritsyan, S. A. Pulbere, S. V. Kotov, A. A. Sokolova, D. D. Pervouchine

## Abstract

Genital lichen sclerosus (GLS) is a chronic inflammatory dermatosis that affects the genital skin. Despite different clinical manifestations, the pathogenesis of GLS in men and women is thought to be common and is attributed to a combination of autoimmune and genetic factors. In this study, we compared the transcriptomic profiles of penile (mGLS) and vulvar lichen sclerosus (VLS) with the objective to identify commonly deregulated genes. We observed a substantial heterogeneity of the transcriptomic signatures in mGLS samples which is driven by different compositions of immune infiltrates. In mGLS, gene expression signatures strongly indicate epidermis dysfunction and overexpression of epithelial inflammation marker Keratin 6 (KRT6) and chitinase CHIT1. No significant changes in the expression levels of known GLS markers such as VIM, CTNNB1, LGALS7 and ECM1 were detected, however, changes in the expression levels of genes associated with autoimmune diseases and genes upregulated in squamous cell carcinoma were observed, including TNF, CCNB1 and RUNX3. There was no enrichment in HCV-derived polyU/UC insertions that were reported previously. Instead, we have identified a long non-coding RNA DRAIC with a large coding potential that is commonly upregulated in mGLS and VLS. Together, our results represent a comprehensive catalog of shared transcriptomic signatures including novel biomarkers and potential therapeutic targets.

## Introduction

Genital lichen sclerosus (GLS) is a chronic inflammatory disease that affects the skin of the genitals in men and women [1]. It manifests itself by white atrophic patches in the affected area, often accompanied by itching, soreness, and painful urination. GLS is characterized by a persistent, recurrent course often leading to severe complications such as cicatricial phimosis, paraphimosis, and urethral stenosis in men [2], and urinary obstruction, vulvar ostium stenosis, and degeneration of genital tissues in women [3]. Furthermore, people of both sexes with GLS are at an increased risk of developing squamous cell carcinoma [4].

In spite of different clinical pictures, the underlying causes of GLS in the two sexes are commonly attributed to a combination of autoimmune and genetic factors, as well as to other conditions such as skin damage or chronic contact with urine [5]. Several molecules have been identified as being associated with the immunopathology of GLS. Among them are the product of the ECM1 gene, a 85-kDa secreted glycoprotein that is expressed in different splice variants, with autoantibodies frequently elevated in GLS patients [6, 7], the miR-155 microRNA known to contribute to sclerotic tissue formation, galectin-7, a pro-apoptotic keratinocyte protein that promotes fibroblast proliferation and others [8, 9]. HLA class II genotypes, particularly HLA-DQ7, are frequently associated with GLS in both sexes, with more than a half of affected females carrying this haplotype [10, 11]. Comorbidity studies indicate that GLS patients frequently have at least one other autoimmune disease [12, 13]. Yet, the molecular causes of GLS are largely unknown, and no universal biomarker currently exists for its definitive diagnosis.

A number of studies approached unveiling the mechanism of GLS pathogenesis by transcriptome profiling such as bulk and single-cell RNA sequencing (RNA-seq) integrated with multi-omics approaches, and also by earlier techniques such as DNA microarrays (Table 1). They generated a number of challenging hypotheses, for instance, on the correlation to abnormal antivirus response due to the presence of Hepatitis C Virus poly U/UC sequences in vulvar lichen sclerosis (VLS) [14], on the host-microbe interactions on the dysbiosis of tissue microbiota in male genital lichen sclerosus (mGLS) [15], and on the role of the crosstalk between fibroblasts and T cells in fibroblast-mediated pathogenesis in the latter [16]. While GLS in males and females is often considered as one etiology [17], their common underlying molecular causes remain unknown and, furthermore, no comparative study of VLS and mGLS has been conducted. In revisiting this problem, we performed high-throughput transcriptome profiling by RNA-Seq of biopsy samples from genital sites of mGLS patients and of healthy donors in order to compare the transcriptomic signatures of VLS and mGLS with the objective of identifying commonly deregulated genes.

**Table 1:** Transcriptome profiling experiments including bulk and single-cell RNA-seq and DNA microarrays.

|  | Bulk RNA-seq | Single cell RNA-seq | MicroArray |
| --- | --- | --- | --- |
| mGLS | Host-microbe interactions on the dysbiosis of tissue microbiota [15] | Fibroblast-mediated pathogenesis [16]; Characteristic subset of cells including fibroblasts [61]; Epigenetic analysis [62] | Gene expression profiling [40]; Gene expression patterns in Congenital phimosis [63] |
| VLS | Transcriptome profiling and network analysis [64]; HCV poly U/UC sequence-induced inflammation [14] | Keratinocytes as key players in the pathogenesis of VLS [46] | Autoimmune phenotype [65] |

## Methods

### Biopsy and sample collection

The clinical study has been approved by the Institutional Review Board of the Skolkovo Institute of Science and Technology and by the Ethics Committee of the N.I. Pirogov Moscow City Clinical Hospital No. 1. In this study, ten male patients between 18 and 42 years of age (median age 28) were enrolled in the Day Hospital of the Urology Division of N.I. Pirogov Moscow City Clinical Hospital No. 1. The GLS group consisted of five patients with a clinically confirmed diagnosis of genital lichen sclerosus. The control group consisted of five healthy individuals undergoing a planned surgical treatment. The operating urologist marked three areas of sclerotically altered foreskin in each patient from the GLS group and three unaffected areas of the foreskin in each patient from the control group, each area comprising a square 0.5 cm x 0.5 cm. Upon dissection, foreskin fragments were minced with a scalpel, transferred into a cryovial, and stored in the liquid nitrogen at -80°C (collection time range from 50 to 85 sec, median collection time 65 sec). The remaining biomaterial was fixed in 10% buffered neutral formalin, compactified and embedded in paraffin. The 4-µm-thick sections of the obtained paraffin blocks were prepared on the microtome, stained with hematoxylin and eosin, and subjected to histological examination.

### RNA extraction

RNA extraction and isolation was done on frozen tissue samples (10-30 mg) using the Purelink mini kit (ThermoFisher, USA). Frozen tissue samples were added to the CK14 tubes precooled to 4°C containing 0.6 ml of the Lysis Buffer (ThermoFisher) and beads (Bertin Technologies). Precellys®24 was applied three times 40 sec at 6300 rpm with 1 min break. Tubes were placed on ice between each cycle. Each sample was centrifuged at 12000 g for 2 min, the extract was moved to a new tube, and 600 µL of 70% ethanol was added. The mixture was vortexed, and 600 µL was transferred to a spin cartridge to centrifuge at 12000 g for 15 sec. The flow-through was discarded, 600 µL of the mixture was added to the spin cartridge again and centrifuged at 12000 g for 15 sec. The cartridge was washed three times by adding 700 µL, 500 µL, and 500 µL of the Wash Buffer I, respectively, followed by centrifuging at 12000 g for 15 sec and discarding the flow-through. The cartridge was placed into a new tube, 100 µL of RNase-free water was added to its center followed by incubation for 1 min and centrifuging at 12000 g for 2 min at room temperature. The RNA concentration was measured by photometry, with the 260/280 nm ratio of optical density in the 2.0-2.1 range. RNA integrity score (RIN) was measured using Agilent 2100 Bioanalyzer on samples containing on average 128 ng/µl RNA yielding the median RIN score of 7. Two samples (mGLS sample 7 and control sample 8) with the RIN score below 6 were discarded. The remaining four mGLS samples and four control samples were chosen for library preparation and RNA-seq.

### RNA-seq libraries and sequencing

PolyA+ RNA libraries were prepared with NEBNext Ultra II Directional RNA Library Prep Kit for Illumina with Purification Beads (E7765 L, New England Biolabs) and NEBNext® Magnetic Bead Poly(A)+ mRNA Isolation Module (E7490 L, New England Biolabs) in accordance to manufacturer protocol. NGS were done and 150-base single-end reads were collected by Illumina 2000.

### cDNA synthesis

One microgram of total RNA was first subjected to RNase-free DNase I digestion (Thermo Fisher Scientific) at 37°C for 30 min to remove contaminating genomic DNA. Next, 500 ng of total RNA were used for complementary DNA (cDNA) synthesis using Magnus First Strand cDNA Synthesis Kit (Evrogene) for reverse transcription-quantitative PCR (RT-qPCR) to a final volume of 20 µl. cDNA was diluted 1:5 with nuclease-free water for quantitative PCR (qPCR).

### RT-qPCR

qPCR reactions were performed in triplicates in a final volume of 12 µl in 96-well plates with 420 nM gene-specific primers and 2 µl of cDNA using 5XqPCRmix-HS SYBR reaction mix (Evrogen). Primers for qPCR are listed in Supplementary Table S1. A sample without reverse transcriptase enzyme was included as a control to verify the absence of genomic DNA contamination. Amplification of the targets was carried out on CFX96 Real-Time System (Bio-Rad), with the following parameters: 95°C for 5 min, followed by 39 cycles at 95°C for 20 s, 60°C for 20 s and 72°C for 20 s, ending at 72°C for 5 min. For each pair of primers in qPCR analysis, the primer efficiencies were estimated using a calibration curve. For each primer pair in the PCR analysis, primer efficiency was assessed using a calibration curve, with primer efficiency exceeding 90% in all cases. Changes in gene expression were calculated using the double normalization method (“ddCt”), taking into account PCR efficiency. The GAPDH gene was used as an internal control to normalize gene expression levels.

### Read Alignment and Transcript Quantification

Adapter trimming and short read filtering were performed using the fastp v0.20 utility [18] with parameters enabling low-quality bases removal ‘-l 35 --cut front --cut right‘. Reads were then aligned to the GRCh38 human reference genome using GENCODE v47 transcriptome annotation with STAR v2.7.8a [19]. The following parameters were used: --outFilterMultimapNmax 20 --alignSJoverhangMin 8 --alignSJDBoverhangMin 1 --outFilterMismatchNmax 999 --outFilterMismatchNoverReadLmax 0.04 --alignIntronMin 20 --alignIntronMax 1000000 -- alignMatesGapMax 1000000 --chimSegmentMin 15. The gene-level counts were generated by assigning the aligned reads to the annotated features using the featureCounts program from the subread package v2.1.1 [20]. Read counts were normalized as TPM using the rnanorm v2.1.0 package. To estimate the fraction of immune cells in a sample, the quantiseq deconvolution method implemented in the immunedeconv package v2.1.0 was used [21, 22].

### Differential gene expression and splicing analysis

Differential gene expression analysis was performed on the raw gene count matrix using the DESeq2 package (pydeseq2 v0.5.0 [23]). In both mGLS and VLS datasets, the GLS samples were compared to tissue samples from healthy donors. The PCA analysis was performed on the VST-transformed gene counts from DESeq. Differential gene expression analysis of TCGA-LUSC paired samples was performed with limma package v3.66 [24]. Genes with the adjusted P-values below 0.05 and the absolute log_2_ fold change greater than one were considered as differentially expressed. The sets of differentially expressed genes were submitted to DAVID web server for functional enrichment analysis [25]. The coding potential of differentially expressed non-coding RNAs was assessed by transdecoder utility v5.7.1. Differential splicing analysis between the disease and control groups was conducted using rMATS v4.3.0 software [26]. The downstream analysis was focused on the exon skipping events, and only the exons with median read coverage of 40 reads in at least one sample group were considered. The events with an adjusted P-value of less than 0.05 and an absolute inclusion level difference greater than 0.05 were considered as differentially spliced.

### Identification of potential neoantigens

The potential presence of microbial and viral sequences was assessed by metagenomic analysis of the RNA-Seq data using Kraken2 v2.1.6 software [27] against the standard database with the default settings. Insertions in reads were quantified using a custom script that extracts the insertion segments from CIGAR strings in BAM files with the corresponding read sequences and alignment coordinates. Insertions containing three or more consecutive uridine nucleotides were counted. The insertions in coding exons were counted separately.

### Statistical procedures

The data were analyzed using python version 3.8.2 and R statistics software version 3.6.3. Benjamini-Hochberg correction was used to account for multiple hypothesis testing and to compute the adjusted *P*-values. Throughout the paper, *r* and *P* denote the Pearson correlation coefficient and the adjusted *P*-value, respectively; *FC* denotes the fold change of gene expression level. Non-parametric tests were performed using normal approximation with continuity correction. In all figures, the significance levels 0.05 and 0.01 are denoted by * and **, respectively.

## Results

### Heterogeneity of transcriptomic signatures in mGLS

High-throughput transcriptome profiling of biopsy samples from the genital skin of four mGLS patients and four healthy donors by RNA-Seq yielded a total of 289 mln. short reads, on average 36 mln. short reads per sample. Gene expression levels were computed from short read counts followed by a normalization by DESeq2 package (Supplementary File 1). Principal Component Analysis (PCA) of gene expression values revealed the lack of clustering by the mGLS vs. control group. Instead, we observed a high degree of heterogeneity within the mGLS cohort, with samples scattering widely along the principal components (Figure 1A). This was particularly evident along the first principal component (PC1) representing 65% of the variance, where samples formed two distinct subgroups, in which samples 1 and 4 were clearly separated from samples 5 and 9. Even when PCA was confined to only differentially expressed genes, the variability of mGLS samples persisted in spite of a clear separation between mGLS and control groups (Figure 1B). The scattering of mGLS samples along the principal components accounting for 80% the variance (PC1 and PC2, 68% and 12%, respectively) suggests the existence of a confounding factor influencing global gene expression profiles.

**Figure 1:**
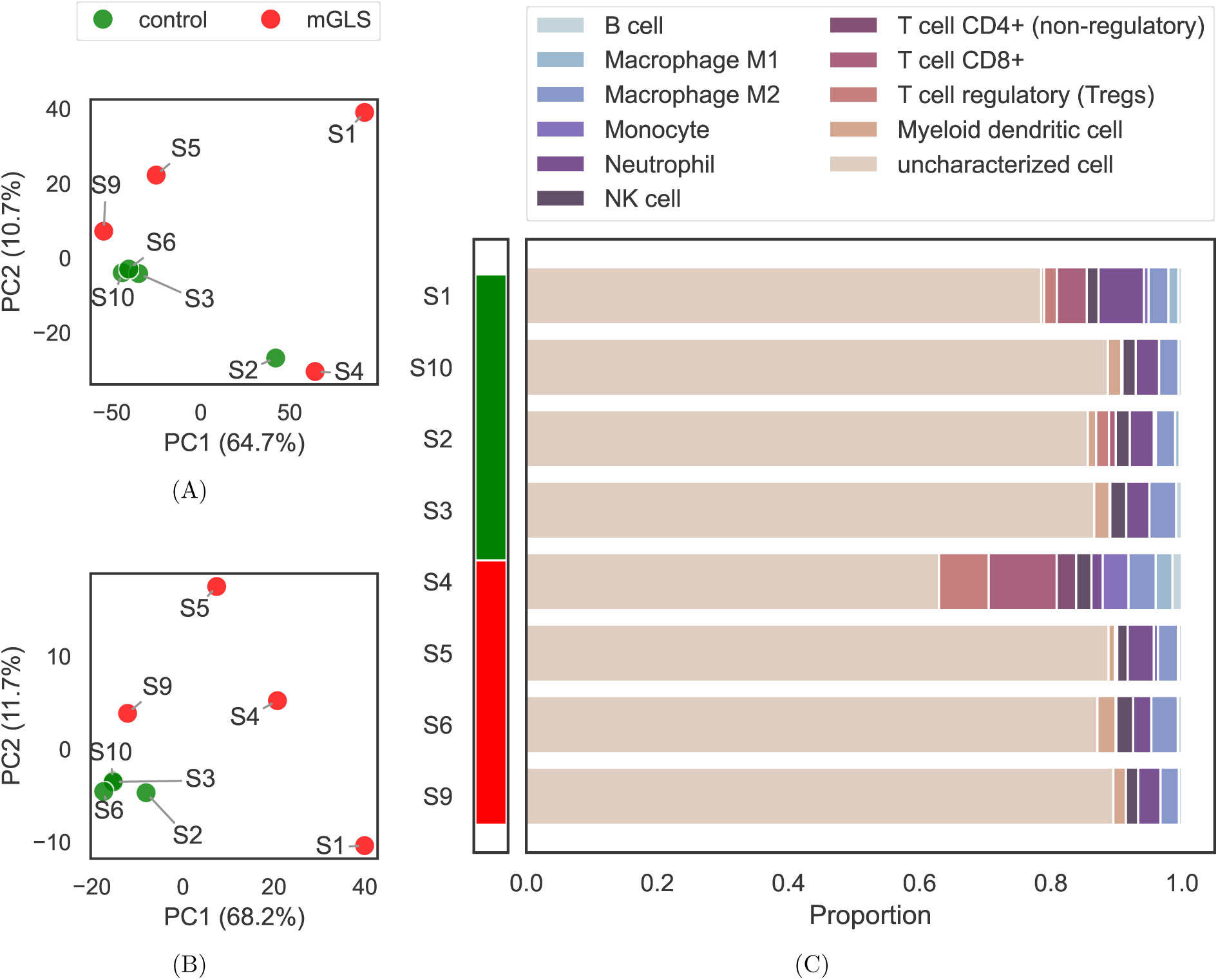
Heterogeneity of transcriptomic signatures in mGLS. **(A)** Principal component analysis (PCA) of expression levels of protein-coding genes (control group: S2, S3, S6, S10; mGLS group: S1, S4, S5, S9). **(B)** PCA with respect to differentially expressed protein-coding genes. **(C)** Proportion of immune cells in mGLS samples estimated by bulk RNA-Seq deconvolution.

Differences in cellular composition, specifically in immune cell infiltration into the diseased tissue, could be a contributing factor to the observed heterogeneity of mGLS samples. To test this hypothesis, we inferred the relative proportions of immune cell subtypes within each sample using a computational deconvolution approach quanTIseq [21]. It revealed a larger and more variable immune cell infiltration in the mGLS group, and only moderate such infiltration in the control group (Figure 1C). Consistent with separate positioning in the PCA plot, samples 1 and 4 exhibited an elevated immune cell proportion and a distinct composition of the immune infiltrate. Sample 4 was characterized by a high relative abundance of CD8+ T-cells (10.4%) and T-regulatory cells (7.6%) suggesting active immune response under immunosuppression [28, 29]. In contrast, immune cell fraction in sample 1 was dominated by neutrophils (6.9%), which may reflect a different inflammatory microenvironment [30]. This divergence from the remaining two mGLS samples, which exhibited relatively low levels of immune cell infiltration likely drives the variation observed in the PCA.

### Gene expression signatures indicate epidermis dysfunction in mGLS

A comparison of transcriptomic profiles of human protein-coding genes using the DESeq2 package has identified 249 differentially expressed genes (DEGs) between mGLS and control samples (*P* < 0.05 and | log_2_ *FC*| > 1). Of these, 164 were upregulated and 85 were downregulated (Figure 2A). Functional enrichment analysis of the upregulated genes using the DAVID tool [31] indicated that the primary transcriptional alteration in mGLS affects epidermal integrity and function. The most significantly enriched Gene Ontology (GO) terms and UniProt keywords (KW) were related to epidermal biology (Figure 2B). The enriched categories included “keratinization” (GO:0031424, *P* < 10^−10^), “cornified envelope” (GO:0001533, *P* = 0.003) and “epidermis” (GO:0030280, *P* = 0.03). The transcriptomic data also contained detectable immune (”innate immunity”, KW-0399, *P* = 0.078) and metabolic signatures (”urea cycle”, KW-0835, *P* = 0.027).

**Figure 2:**
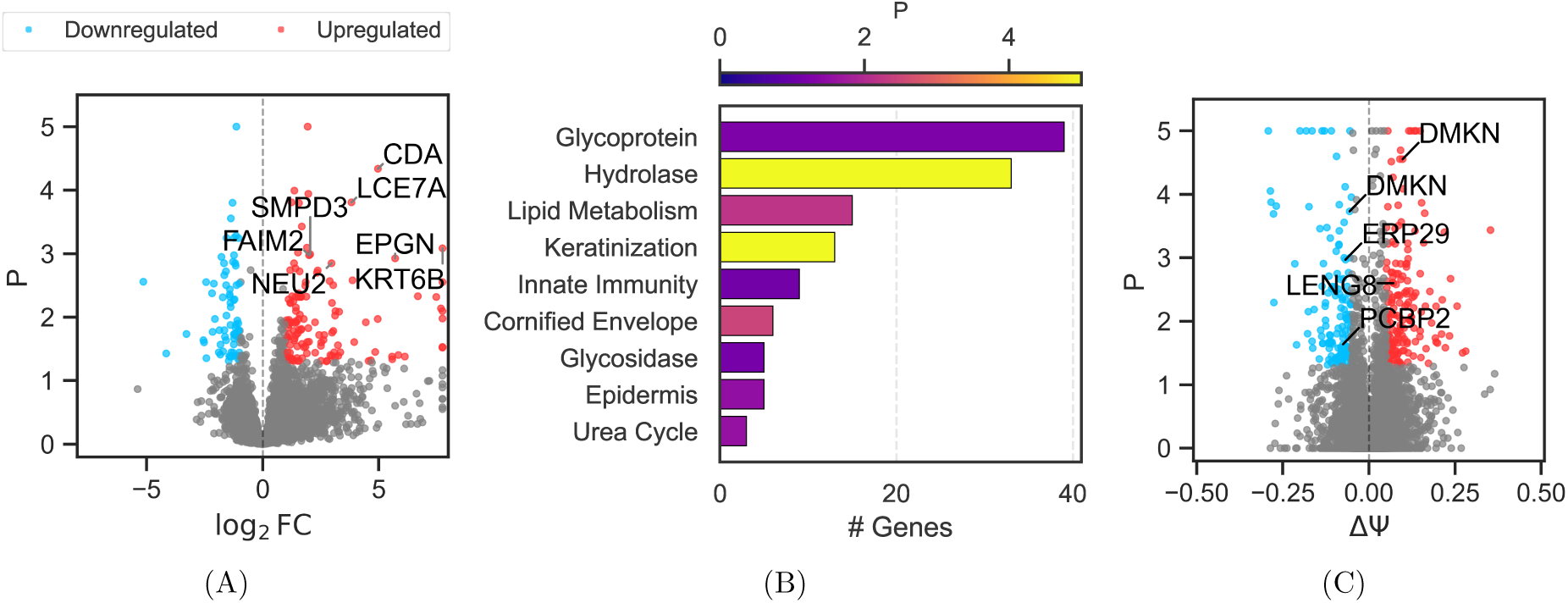
mGLS transcriptomic profile. **(A)** The volcano plot of gene expression. Significantly deregulated genes (*P* < 0.05 and | log_2_ *FC*| > 1) are shown in color. **(B)** Gene set enrichment analysis of upregulated genes. Gene Ontology (GO) categories with *P* < 0.1 are shown. **(C)** The volcano plot of differentially spliced cassette exons. Significantly deregulated events (*P* < 0.05 and |ΔT| > 0.05) are shown in color.

In addition to the analysis of gene expression signatures, we performed differential analysis of alternative splicing by estimating exon inclusion ratios (T, PSI, percent-spiced-in) defined as the number of split reads supporting exon inclusion as a fraction of the combined number of split reads supporting exon inclusion and skipping. A number of exons underwent significant splicing changes in mGLS as evidenced by the change in exon inclusion ratio (ΔT), remarkably in some genes associated with autoimmune disorders (Figure 2C). Among them are two exons in the dermokine (DMKN) gene, which encodes a skin-specific secreted glycoprotein [32, 33] implicated in inflammatory bowel disease [34], an exon in the leukocyte receptor gene (LENG8), and an exon in the poly(C)-binding protein PCBP2, both of which play a role in regulating T-cell function [35, 36]. All differentially spliced exons are listed in Supplementary File 2.

### Shared transcriptomic signatures in mGLS and VLS

To reveal the common molecular mechanisms underlying the pathophysiology of GLS in the two sexes, we compared the transcriptomic profiles obtained for the male cohort with the transcriptomic profiles of VLS from a publicly available dataset [14]. The female cohort consisted of nine matched pairs (tissues affected by VLS and adjacent healthy tissues) from the same donors, and an additional group from four healthy donors. To ensure a comparable analysis, we identified DEG harbored in autosomes by comparing patient-derived VLS samples against samples from healthy donors. This combined dataset allowed for a cross-sex comparison of the GLS transcriptomes.

The comparison conducted separately in the two sexes revealed a notable disparity in the scale of transcriptomic changes. In the male cohort, we identified 249 DEGs, with 164 being upregulated and 85 being downregulated. By contrast, the female cohort exhibited a more extensive deregulation, with 1116 significantly upregulated and 1112 downregulated genes. This difference in both the magnitude and the balance of DEGs may be attributable to the larger size and hence higher statistical power in the female dataset, as well as to biological differences such as larger and compositionally distinct immune infiltrate in the male cohort.

The expression values of protein-coding genes (Figure 3A) showed a weak but significant positive correlation between mGLS and VLS (*r* = 0.21, *P* < 10^−10^) indicating that the transcriptional changes in these diseases are not identical, but pathogenic alterations or their downstream effects may be similar. One of consistently upregulated genes in both mGLS and VLS was Keratin 6 (KRT6), a marker of hyperproliferative and wounded skin states such as psoriasis [37]. Its shared overexpression aligns with the characteristic lichen sclerosus manifestations of lesion formation. A shared overexpression was also detected for the CHIT1 gene, which codes for the enzyme chitinase 1 and is not directly linked to GLS, however other members of the chitinase family are known as biomarkers in immune-mediated diseases [38].

**Figure 3:**
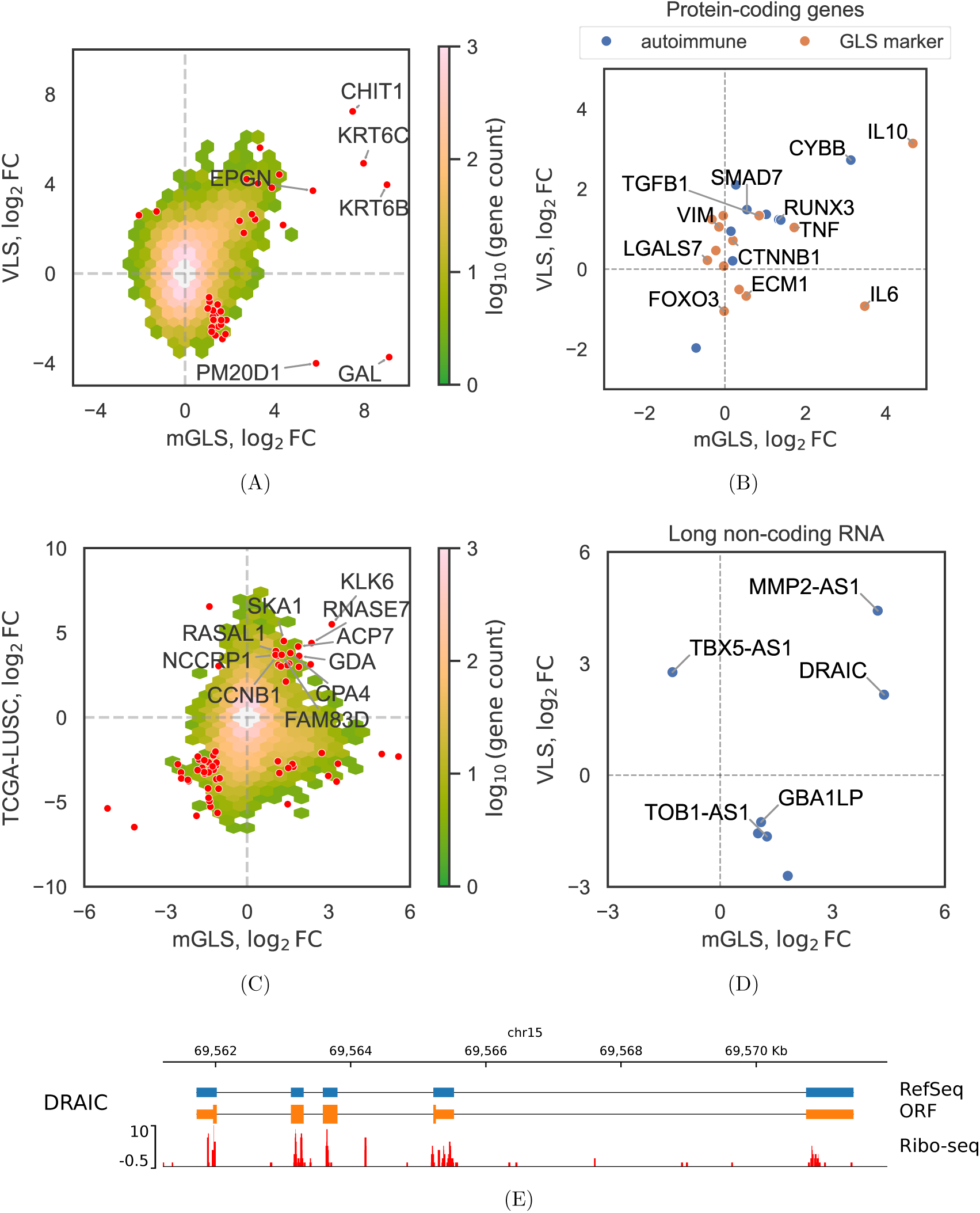
Shared transcriptomic signatures in mGLS and VLS. **(A)** Density plot of log_2_ protein-coding gene expression fold changes (log_2_ *FC*, GLS vs. control) in mGLS and VLS. The colormap represents log_1_ 0 of the number of genes in each bin. Genes significantly deregulated (*P* < 0.05 and | log_2_ *FC*| > 1) in both mGLS and VLS are indicated by red dots. **(B)** Expression levels of protein-coding genes with known links to GLS or autoimmune diseases (Table 2). **(C)** Density plot of protein-coding gene expression fold changes in mGLS and TCGA-LUSC (GLS vs. control and tumor vs. control, respectively). Genes significantly deregulated in both datasets are indicated by red dots. **(D)** Same as (A) for long non-coding RNAs (lncRNA). **(E)** Genomic organization of the DRAIC lncRNA including the annotated transcript (RefSeq), the predicted open reading frame (ORF), and aggregated Ribo-seq signal from GWIPS-viz (Ribo-seq).

**Table 2:** log_2_FC values for genes with known links to GLS or autoimmune diseases.

| ENSEMBL gene ID | Gene name | Description | mGLS | VLS |
| --- | --- | --- | --- | --- |
| ENSG00000108821 | COL1A1 | Collagen type I alpha 1 chain | -0.04 | 1.33 |
| ENSG00000168542 | COL3A1 | Collagen type III alpha 1 chain | -0.32 | 1.24 |
| ENSG00000130635 | COL5A1 | Collagen type V alpha 1 chain | -0.23 | 0.46 |
| ENSG00000143369 | ECM1 | Extracellular matrix protein 1 | 0.53 | -0.67 |
| ENSG00000118689 | FOXO3 | Forkhead box O3 | -0.02 | -1.05 |
| ENSG00000136634 | IL10 | Interleukin 10 | 4.67 | 3.14 |
| ENSG00000136244 | IL6 | Interleukin 6 | 3.47 | -0.93 |
| ENSG00000178934 | LGALS7B | Galectin 7B | 0.35 | -0.51 |
| ENSG00000105329 | TGFB1 | Transforming growth factor $\beta$ 1 | 0.85 | 1.34 |
| ENSG00000232810 | TNF | Tumor necrosis factor | 1.72 | 1.04 |
| ENSG00000141510 | TP53 | Tumor protein p53 | -0.03 | 0.08 |
| ENSG00000168036 | CTNNB1 | Catenin $\beta$ 1 | 0.20 | 0.72 |
| ENSG00000026025 | VIM | Vimentin | -0.15 | 1.05 |
| ENSG00000165168 | CYBB | Cytochrome b-245 β-chain | 3.12 | 2.72 |
| ENSG00000120738 | EGR1 | Early growth response 1 | -0.72 | -1.97 |
| ENSG00000179348 | GATA2 | GATA binding protein 2 | 0.15 | 0.95 |
| ENSG00000183019 | MCEMP1 | Mast cell expressed membrane protein 1 | 0.27 | 2.10 |
| ENSG00000105835 | NAMPT | Nicotinamide phosphoribosyltransferase | 0.19 | 0.21 |
| ENSG00000020633 | RUNX3 | RUNX family transcription factor 3 | 1.34 | 1.24 |
| ENSG00000143546 | S100A8 | S100 calcium binding protein A8 | 1.03 | 1.37 |
| ENSG00000163220 | S100A9 | S100 calcium binding protein A9 | 1.39 | 1.23 |
| ENSG00000101665 | SMAD7 | SMAD family member 7 | 0.55 | 1.49 |

Next, we selected genes with known links to GLS or autoimmune diseases (Table 2) including vimentin (VIM) and beta-catenin (CTNNB1), two genes previously identified as histological markers in mGLS, thirteen genes involved in GLS pathogenesis as listed in a prior review [17], and nine genes known to be deregulated in at least two autoimmune conditions [39] (Figure 3B). Contrary to expectations, the levels VIM and CTNNB1 were not elevated at the transcript level in mGLS, while in VLS their expression levels slightly increased. Within the set of thirteen GLS-associated genes, we examined two key candidates: the proposed autoantigen ECM1 that was previously reported as downregulated specifically in mGLS [40] and galectin-7 (LGALS7), but neither of them showed a significant deregulation. However, several immunity-related genes were significantly altered. Tumor necrosis factor-alpha (TNF) was significantly upregulated in both mGLS and VLS, while interleukin-6 (IL-6) was upregulated specifically in the male patients, but suppressed in VLS patients. The elevation of these proinflammatory cytokines is consistent with the activation of inflammatory response in GLS. For the group of genes associated with the progression of autoimmune diseases, we found a significant upregulation of cytochrome B β-chain (CYBB), SMAD Family Member 7 (SMAD7), and RUNX Family Transcription Factor 3 (RUNX3) in both mGLS and VLS.

### mGLS and squamous cell carcinoma

It has been known that mGLS is associated with an increased risk of malignant transformation [41]. To characterize the molecular basis of this association, we compared mGLS gene expression profiles with those of lung squamous cell carcinoma (LUSC) from The Cancer Genome Atlas (TCGA) in the absence of an available transcriptomic dataset for penile carcinoma. We identified 17 genes that were commonly upregulated in both mGLS dataset and in the paired samples of TCGA-LUSC cohort (Figure 3C). The Gene Ontology (GO) analysis of this shared gene set demonstrated an enrichment for genes associated with cell division (”cell division”, KW-0835, *P* = 0.0084). Among the shared upregulated genes were the G2/mitotic-specific cyclin-B1 (CCNB1) and Ribonuclease A Family Member 7 (RNASE7), which was elevated by a factor of 2.14 in mGLS and by a factor of 12.38 in LUSC. The gene encoding RNASE7 was also consistently upregulated by a factor of 5.17 in mGLS cohort and by a factor of 20.97 in LUSC.

### Neo-antigens and long non-coding RNAs

A leading hypothesis about the pathogenesis of autoimmune diseases is the presentation of foreign or altered self-antigens that trigger aberrant immune response. Earlier works have identified the ECM1 protein as a potential autoantigen and showed the presence of anti-ECM1 antibodies in the majority of patients [7]. However, subsequent studies suggested that ECM1 autoantibodies are not involved in triggering the onset of GLS and instead represent an epiphenomenon of the disease progression [42]. We therefore focused on characterizing pathogenic or endogenous transcripts with a potential to generate novel immunogenic peptides.

An earlier study of VLS by RNA-Seq proposed that the autoimmune response could be triggered by the Hepatitis C virus (HCV) polyU/UC motif integrated into the host genome [14]. While the mechanism of such integration remains elusive for an RNA virus without reverse transcription activity, evidence exists for the presence of HCV sequences in the host DNA [43]. In revisiting this hypothesis, we performed a direct taxonomic classification of short read sources using Kraken2 [27] on both male and female GLS datasets. It revealed that viral reads constitute less than 1% of all reads with no detectable enrichment in GLS samples. Furthermore, we assessed the potential for altering protein-coding sequences by counting the genomic and exonic insertions in RNA-Seq read alignments that contain three or more consecutive uridine nucleotides. No enrichment for such insertions was observed in GLS patients, and the frequency of exonic insertions was remarkably low, on the order of one hundred reads per sample (Supplementary Figure 1). Overall, our analysis does not support the presence of HCV-derived polyU/UC sequences in GLS transcriptomes. The original association [14] may have originated from misaligning short reads with polyA tails to viral genomes, which generates false-positive signals for polyU/UC-like motifs.

We subsequently checked endogenous sources of neoantigens among long non-coding RNAs (lncRNAs) that contain unannotated open reading frames with a potential to encode novel peptides. Differential gene expression analysis identified a total of 7 lncRNAs with significant changes in the expression level (*P* < 0.05 and | log_2_ *FC*| > 1 in both datasets), and only two of them were upregulated in both male and female patients (Figure 3D). One of them, the DRAIC lncRNA was found to contain a candidate open reading frame located on its strand (Figure 3E). The translational potential of DRAIC reflected by Ribo-seq data from the GWIPS-viz portal [44] showed a weak but consistent signal in its genomic locus. Another transcript commonly overexpressed in both mGLS and VLS is MMP2 Antisense RNA 1 (MMP2-AS1), which is associated with lung non-small cell cancers [45]. However the expression levels of both transcripts were remarkably low (TPM=0.18 and 0.78, respectively). Other non-coding RNA species didn’t show a concordant and significant change in mGLS and VLS.

### Validation of deregulated genes

In order to validate deregulation of markers predicted from RNA-seq experiments, we performed RT-qPCR analysis of the selected genes in five technical replicates for RNA obtained from each of the eight patients. The gene expression level measured by RT-qPCR was normalized to that of the glyceraldehyde-3-phosphate dehydrogenase (GAPDH) gene, which was used as a reference. As evidenced by FC values with respect to the median value in the control group, KRT6C, TNF, CCNB1, and RNASE7 were significantly overexpressed in mGLS samples compared to the control samples, with a remarkable number of outliers presumably reflecting transcriptomic heterogeneity (Figure 4). The expression levels of VIM and TGFB1 were not significantly different between mGLS and control groups, while the expression level of DRAIC lncRNA was at borderline detection level in both, consistent with its low TPM counts in RNA-seq data. The latter result doesn’t invalidate its role in the pathogenesis of mGLS, since even low-expressed transcripts can produce immunogenic peptides, which may have a significant impact on the immune response.

**Figure 4:**
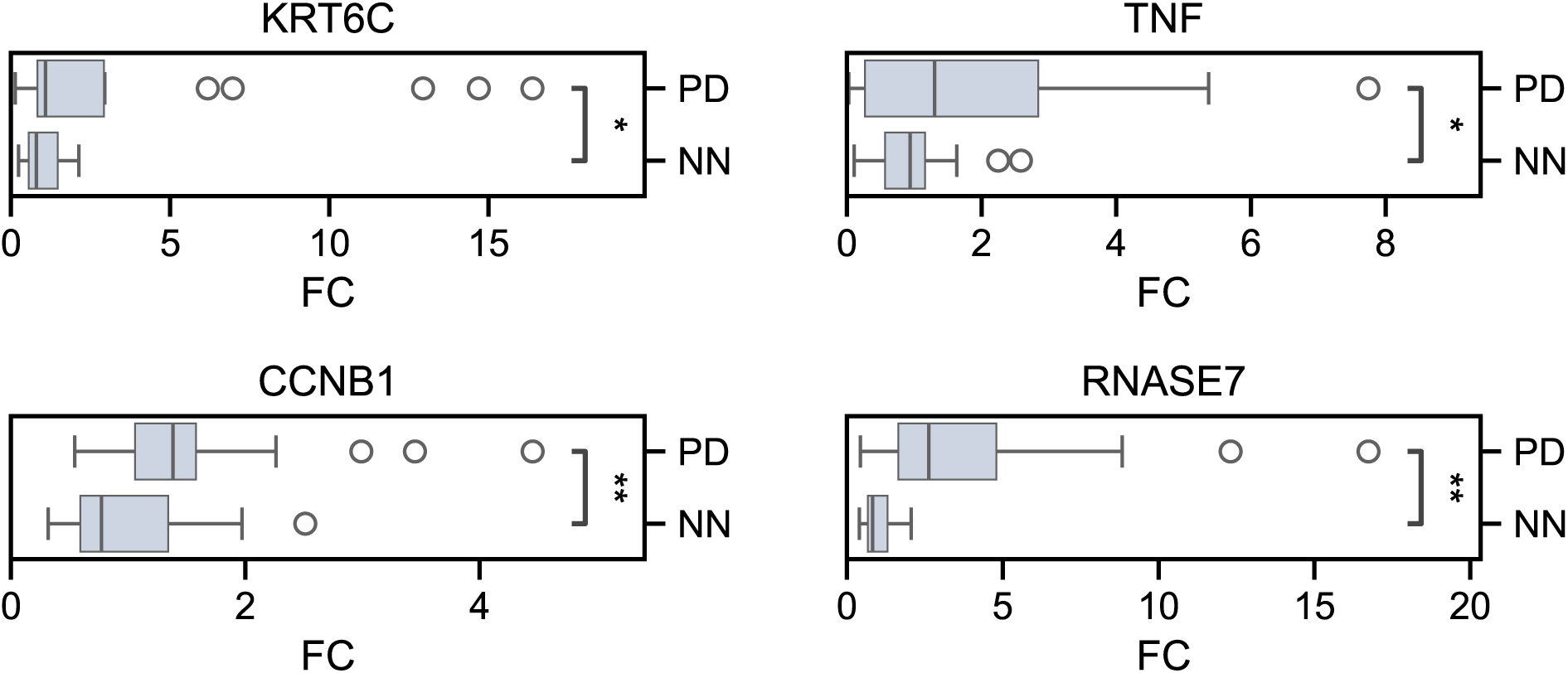
RT-qPCR validation of upregulation for Keratin 6C (KRT6C), tumor necrosis factor (TNF), Cyclin B1 (CCNB1) and Ribonuclease A Family Member 7 (RNASE7). Gene expression levels were normalized by the expression levels of GAPDH. PD – mGLS group. NN – control group. Boxplots represent expression fold change (FC) relative to the mean expression in the control group. For each gene, 5 independent replicates were examined in each of the 8 patients. Statistically significant differences at the significance levels 5% and 10% are denoted by * and **, respectively.

## Discussion

Heterogeneity of transcriptomic signatures inferred from RNA-seq experiments is often attributed to different composition of cellular subtypes comprising tissue samples, although other factors such as variation in the presentation, clinical manifestations, and severity of the disease also plays a key role. Histological studies have demonstrated a large interpatient heterogeneity in lesional skin from VLS patients [46]. Transcriptomic heterogeneity in mGLS has also been reported, with two distinct subtypes corresponding to the activation of immune and epithelial cell proliferation pathways, respectively, while sharing common signatures related to hyperkeratosis [47]. In our study, the observed heterogeneity of the transcriptomic signatures in mGLS samples can primarily be attributed to a different composition of the immune infiltrate, namely in the relative abundance of CD8+ T-cells, T-regulatory cells, and neutrophils, which is responsible for the lack of clustering of mGLS samples in the PCA plot. More detailed estimates of GLS heterogeneity can be inferred by analyzing single-cell RNA-seq experiments (Table 1), which falls outside the scope of this short report.

Remarkably, the gene set enrichment analysis has identified the most overrepresented functional gene categories related to the integrity of epidermal cutaneous structure and innate immunity, but also to metabolic categories such as urea cycle. The latter was driven by the upregulation of three genes including two arginases (ARG1, ARG2), which could have implications on immunity, as the arginine catabolism by myeloid cells is known to contribute to the modulation of immune response [48]. However, in spite of these immune-related signals, the change in epidermal transcriptional programs was the most pronounced alteration. Functional alternations in splicing programs are harder to assess since the exact roles of splice isoforms are largely unknown. Nevertheless, it appears plausible that splicing changes in the DMKN gene produced by keratinocytes could reflect the shift to DMKN-β splice isoform in response to proinflammatory cytokines [49].

The lack of the expected upregulation of known GLS markers such as VIM and CTNNB1 in both mGLS and VLS suggests a large discrepancy between protein-level histology and mRNA abundance. For ECM1, the lack of deregulation aligns with the previous study, which reported this effect only in pediatric-onset cases but not in the adult-onset cohort [17]. The absence of LGALS7 deregulation, which has been implicated in fibroblast proliferation in VLS [50], may again be attributed to the differences between immunohistochemical staining without PCR confirmation in whole-tissue samples [50]. A number of genes associated with the progression of autoimmune diseases were activated in both mGLS and VLS, including CYBB, SMAD7 and RUNX3. They play distinct roles in immune regulation: CYBB is important for reactive oxygen species production in phagocytes [51], SMAD7 inhibits TGF-β signaling by promoting

TGF-β degradation [52], which often results in pro-inflammatory effect [53], and RUNX3 also modulates TGF-β responses in dendritic cells [54]. Their upregulation supports the involvement of immune response in GLS pathogenesis consistent with the hypothesis of its autoimmune origin, however without any clear mechanistic model [39].

It is remarkable that two non-coding transcripts (DRAIC and MMP2-AS1) that were up-regulated in both mGLS and VLS are related to cancer pathogenesis. DRAIC is known to function as a tumor suppressor in prostate and other cancers by inhibiting NF-kB signaling [55]. It was not previously reported as a protein-coding gene, however Ribo-seq experiments weakly indicate that DRAIC may actually be translated. The MMP2-AS1 lncRNA is known to contribute to the progression of renal cell carcinoma by modulating miR-34c-5p/MMP2 axis [56]. Despite the absence of open reading frames in MMP2-AS1, antisense transcripts often encode micropeptides with important functions in cancer [57]. Genes commonly upregulated in mGLS and in non-small cell cancers included CCNB1, a central regulator of the G2/M phase transition [58] which is upregulated in many types of cancer and linked to poor prognosis [59], and RNASE7, an antimicrobial protein secreted by various epithelial tissues and linked to cutaneous squamous cell carcinoma [60]. All these mRNAs could be considered as common potential biomarkers of GLS or targeted by therapeutic modulatory approaches.

## Conclusion

Genital lichen sclerosus has long been considered an enigmatic and challenging disease. Here we conducted a comparative survey of transcriptomic signatures, which revealed multiple genes that are commonly deregulated in mGLS and VLS, including previously unreported biomarkers KRT6 and CHIT1, genes with known links to GLS and autoimmune diseases (TNF, CYBB, SMAD7, RUNX3), long non-coding RNAs DRAIC and MMP2-AS1, as well as genes commonly deregulated in GLS and in squamous cell carcinomas. These results add to the bulk of the knowledge about GLS pathogenesis and pave the road to its correct diagnosis and treatment.

## Authors’ contributions

D.D.P and A.A.S. designed the study; E.M.A., M.M.I., S.A.P., S.V.K. and A.A.S. conducted the clinical part and sample collection, A.L.K. and D.A.S. performed sample processing and library preparation, S.D.M and D.D.P performed data analysis; S.D.M. and D.D.P. wrote the first draft of the manuscript. All authors edited the final version of the manuscript.

## Acknowledgments

The authors thank Elena Shagimardanova and Genomics Core Facility of Skolkovo Institute of Science and Technology for help with high throughput RNA sequencing experiments.

## LIST OF ABBREVIATIONS

DEGs: differentially expressed genes
RT-PCR: quantitative reverse transcription PCR
GLS: genital lichen sclerosus
mGLS: male genital lichen sclerosus
VLS: vulvar lichen sclerosus
PCA: principal component analysis
TMP: transcripts per million.

