## Supplementary material for "A comparative study of genital lichen sclerosus transcriptomes"

### SUPPLEMENTARY INFORMATION: Shared transcriptomic signatures of male and female genital lichen sclerosus

December 10, 2025

### List of Figures

### List of Tables

S1 Primers for RT-qPCR. . . . . . . . . . . . . . . . . . . . . . . . . . . . . . 4

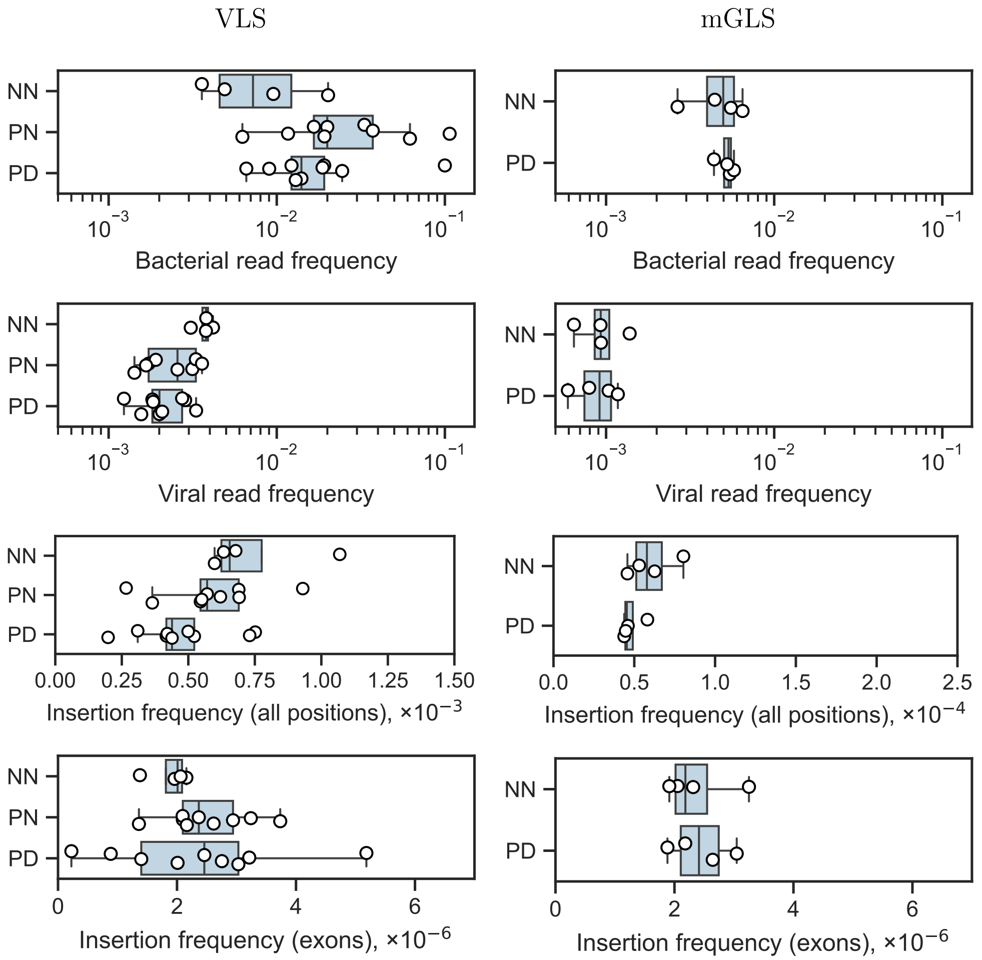


**Figure S1:** Potential sources of neoantigens in mGLS and VLS. The frequency of microbial reads (bacterial and viral) detected by Kraken2, as well as the frequency of polyuridine insertions (genomic and exonic) are shown.

| Primer | Sequence |
|---|---|
| GAPDH_F | GTCTCCTCTGACTTCAACAGCG |
| GAPDH_R | ACCACCCTGTTGCTGTAGCCAA |
| KRT6C_F | ATGGACAACAACCGCAACCT |
| KRT6C_R | ATGTCTGCCTGCTGTGACC |
| CCNB1_F | CGGCCTCTACCTTTGCACTT |
| CCNB1_R | GGCCAAAGTATGTTGCTCGAC |
| TNF_F | TGCACTTTGGAGTGATCGGC |
| TNF_R | GGCCAGAGGGCTGATTAGA |
| RNASE7_F | AGAGATCCCAGTCAGTGCCA |
| RNASE7_R | GGGCTGCATGTGCTGAATTT |

**Table S1:** Primers for RT-qPCR.
